# Cdc48 regulates intranuclear quality control sequestration of the Hsh155 splicing factor

**DOI:** 10.1101/2020.06.16.152934

**Authors:** Veena Mathew, Arun Kumar, Yangyang K. Jiang, Kyra West, Annie S. Tam, Peter C. Stirling

## Abstract

Cdc48/VCP is a highly conserved ATPase chaperone that plays an essential role in the assembly or disassembly of protein-DNA complexes and in degradation of misfolded proteins. We find that Cdc48 accumulates during cellular stress at intranuclear protein quality control (INQ) sites. Cdc48 function is required to suppress INQ formation under non-stress conditions and to promote recovery following genotoxic stress. Cdc48 physically associates with the INQ substrate and splicing factor Hsh155 and regulates its assembly with partner proteins. Accordingly, *cdc48* mutants have defects in splicing and show spontaneous distribution of Hsh155 to INQ aggregates where it is stabilized. Overall, this study shows that Cdc48 regulates deposition of proteins at INQ and suggests a previously unknown role for Cdc48 in the regulation or stability of splicing subcomplexes.

## Introduction

Protein quality control (PQC) refers to the process of triage for non-native, unassembled or mis-localized proteins through a combination of sequestration, degradation and refolding (1). PQC is a part of normal cellular homeostasis and is particularly important during stress when non-native proteins assemble in visible aggregate structures at different subcellular locations. Various proteotoxic stresses elicit such a coordinated PQC response, with the best characterized being heat stress (2), or expression of amyloid-forming proteins(3). PQC responses depend on networks of molecular chaperones, on the action of stress activated protein aggregase and disaggregase activities (1), and on protein degradation by the ubiquitin-proteasome system and autophagy (3, 4).

During stress in yeast, specific types of aggregate structures with different components and behaviors form (5). In the cytoplasm, aggregates at the vacuolar membrane called the Insoluble Protein Deposit (IPOD) are repositories for irreversibly aggregated proteins destined for degradation, such as amyloids (3). Smaller dynamic cytoplasmic aggregates called Q-bodies (quality control bodies) have also been described for protein aggregate reporters such as Ubc9-ts and VHL (6-8). Finally, at the nuclear envelope the juxtanuclear quality control site (JUNQ) forms peripheral to but outside the nucleus, while the intranuclear quality control site (INQ) forms peripheral to and inside the nucleus(3, 9). Most yeast studies of PQC focus on aggregation prone reporter proteins, such as poly-glutamine fragments, temperature-sensitive (ts) mutants or misfolding human proteins (3, 6, 7, 9). Recently, endogenous proteins have been used as markers of aggregate formation, in particular following genotoxic stress (10-12). Following DNA damage, specific nuclear proteins are actively sequestered in INQ structures (10, 11). Focusing on different INQ substrates, it has been suggested that INQ sequestration helps to regulate the replication fork checkpoint (i.e. Mrc1(10)), the activation of DNA repair proteins (i.e. Mus81 (12)), or rewiring of the spliced transcriptome (i.e. Hsh155 (11)).

INQ formation is driven by the small heat shock protein Hsp42, and the aggregase protein Btn2 (13), while INQ dissolution is regulated by Hsp104 and the Hsp70 system(9, 11, 12). Here we show that Cdc48 is also a stress-inducible component on INQ structures and that it plays an enzymatic role in INQ turnover. Cdc48 plays numerous normal and stress responsive roles in cells, including in nuclear PQC (14-17). Here we show that Cdc48 is concentrated at INQ following DNA damage, and that its activity is important to suppress spontaneous aggregation of splicing factor Hsh155 in INQ. Cdc48 physically interacts with Hsh155 even under non-stress conditions and promotes its assembly with partners in the SF3B complex of the spliceosome. Together these results demonstrate a previously unrecognized role for Cdc48 in protein triage to the INQ and suggest unappreciated roles for Cdc48 in splicing.

## Results and Discussion

### Cdc48-GFP localizes to intranuclear quality control sites

We previously screened hundreds of genome stability regulatory proteins for relocalization in response to DNA damage, and identified uncharacterized stress-induced relocalization of Cdc48-GFP (11). Cdc48 is widely distributed in unstressed cells, but is enriched at a nuclear periphery/ER-localization due to its role in ER-associated degradation (ERAD) (**Figure 1A**). Following methyl methanesulfonate (MMS) treatment, Cdc48 relocalized to nuclear and non-nuclear foci (**Figure 1A**). To test if these foci were INQ, we expressed Cdc48-GFP together with Hos2-mCherry, a known marker of the INQ (10, 18). Cdc48 foci colocalized with Hos2-mCherry foci, suggesting that Cdc48 was accumulating at INQ following MMS treatment (**Figure 1B**). Cdc48 also colocalized with Hos2 at peripheral aggregates, which may be cytoplasmic PQC (CytoQ) sites (**Figure 1B and C**). We next ablated INQ formation by deleting the chaperones *HSP42* and *BTN2*, which are essential for INQ protein localization (9-11). Cdc48-GFP foci were strongly diminished in MMS-treated *hsp42*Δ or *btn2*Δ cells, showing that INQ formation was essential for Cdc48 foci (**Figure 1D**). In contrast, deletion of the Hsp40 homologue Apj1, which was recently implicated in disaggregation and turnover of INQ resident proteins, dramatically increased the frequency of Cdc48-GFP foci (**Figure 1D**) (19). Together these experiments suggest that Cdc48 foci mark INQ structures induced by MMS treatment and are influenced by INQ regulatory chaperones.

**Figure 1.**
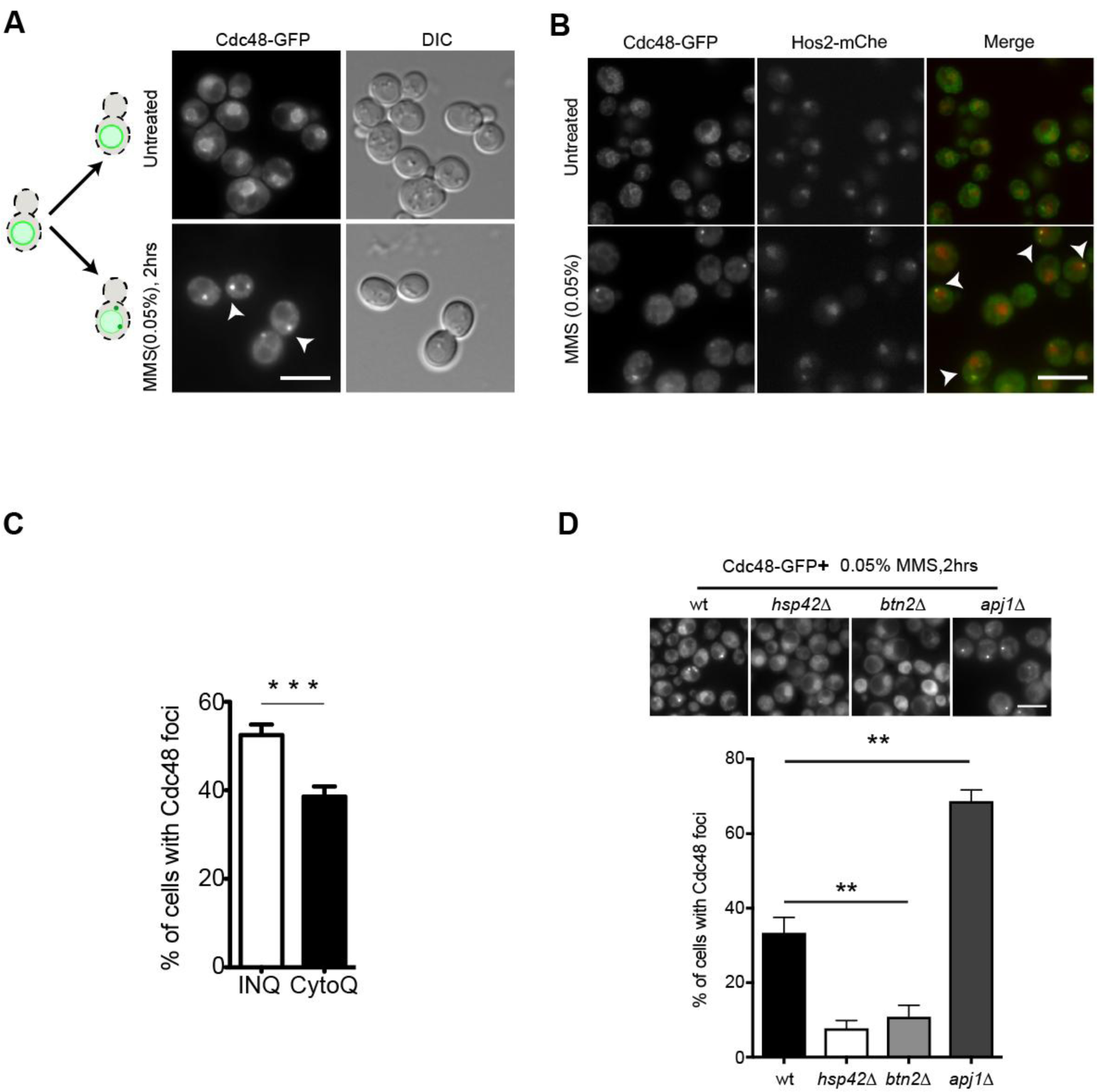
Cdc48 localizes to PQC in a chaperone dependent manner. (A) Cdc48-GFP fusions relocalize into foci upon MMS treatment (white arrowhead). A schematic on left summarizes the movements in both conditions. (B) Cdc48-GFP foci overlap (white arrowheads in merge) with Hos2-mCherry (mChe) foci after MMS treatment. (C) Quantification of MMS induced Cdc48 in perinuclear (INQ) and peripheral (CytoQ) foci. Mean ± SEM, n= 3 with >100 cells each. (Asterisks show *p*-value (***) p<0.001, Fisher test). (D) Reduction of Cdc48-GFP foci in cells lacking *HSP42* or *BTN2* and increase in *apj1*Δ cells. Representative images (above) and quantification of percentage of cells with foci (below). **indicates p<0.01. Error bars are mean± SEM, n= 3 with >50 cells each. Fisher test. Asterisks show *p*-value thresholds in comparison to WT under the same condition. Scale bar, 5μm and for all subsequent figures.

### Cdc48 ATPase regulates the aggregation and INQ localization of Hsh155

Since Cdc48 functions in protein biogenesis and turnover, we suspected that it might have a functional role at INQ. We assembled a set of *CDC48* ts-alleles, which grew robustly at the permissive temperature of 25°C but exhibited a range of fitness defects (**Fig. S1A**) (20). Examining the localization of an INQ substrate in the most severe allele, *cdc48-4601* (hereafter called *cdc48-4*) (**Fig. S1C**), revealed spontaneous Hsh155-GFP and Hos2-GFP foci at nuclear and cytoplasmic aggregate sites (**Figure 2A** and **Fig. S1B**). MMS treatment further increased the proportion of cells with visible aggregates beyond WT cells (**Figure 2B**). We confirmed that other *CDC48* ts-alleles also increased spontaneous INQ formation to varying degrees (**Figure 2C**). Consistent with a functionally important role, *cdc48-4* alleles were significantly impaired in recovery from a two-hour treatment of MMS (**Figure 2D**). While there are many possible mechanisms for the MMS sensitivity of a *cdc48-ts* allele, our data show that Cdc48 function is important for regulating the INQ localization of Hsh155 in normal and DNA damaging conditions. Finally, since Cdc48 uses its ATPase activity to disassemble or unfold proteins, we tested the ability of enzymatically active or dead alleles to rescue the *cdc48-4* phenotype. While WT *CDC48* plasmids suppressed Hsh155 foci, an ATPase-deficient *cdc48*^*E315Q*^ allele could not (**Figure 2E** and **F**) (16). Thus, the Cdc48 ATPase activity is essential to suppress aggregation of Hsh155-GFP into INQ structures.

**Figure 2.**
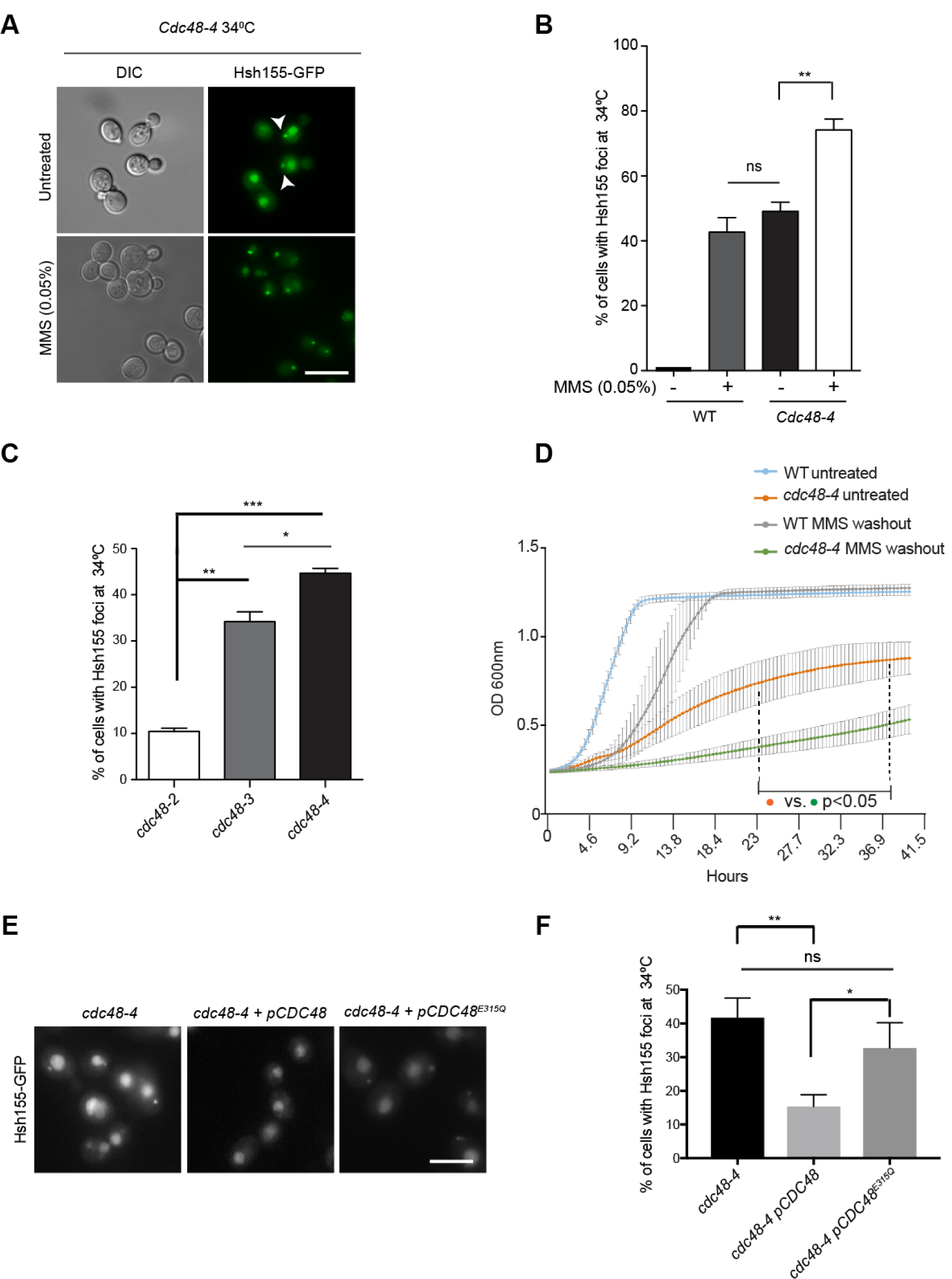
Cdc48 ATPase regulates the aggregation and INQ localization of Hsh155. (A) Visualization of Hsh155-GFP foci in cells with temperature sensitive (ts) alleles of *CDC48* with or without MMS stress at 34°C. White arrowheads show foci in untreated cells. (B) Quantification of foci accumulation in untreated and MMS treated *cdc48-4* strains compared to wild type (WT) strains with or without MMS. (C) Spontaneous Hsh155-GFP foci in three different *CDC48* mutant alleles at 34°C. (D) Growth curve analysis of *cdc48-4* after 2hrs of MMS treatment and washout compared to WT strains ± MMS washout at 34°C. The dotted line represents the range wherein all data points significantly differ from WT after MMS washout. N=3 with triplicates in each. Error bars represent mean± SEM, two way ANOVA with Tukey’s posthoc test, p-value ranges from <0.05- <0.0001. Significant differences between untreated and treated cells are indicated in the interval, *p<0.05. (E and F) Rescue of Hsh155-GFP foci formation by *CDC48-wt* but not *CDC48*^*E315Q*^ ATPase-deficient allele. E is a representative image of data quantified in F. For B, C and F quantifications: Error bars are mean± SEM, n= 3 with >50 cells each. Fisher test, Asterisks show *p*-value thresholds (*** p<0.001; ** p<0.01; * p<0.05 and non-significant (ns)).

### SUMOylation regulates INQ formation in cdc48 mutants

Previous studies link Cdc48 and its co-factors to proteostasis of nuclear proteins via the San1 E3 ligase, and the inner nuclear membrane proteins Asi1-Asi3, however we see no significant effect of *SAN1* deletion on INQ (**Fig. S1D**) (16, 17). In addition, Cdc48 regulates protein-DNA complexes, such as the removal of RNA polymerase from damaged DNA (15), the removal of replisomes during DNA replication termination (14), at protein-DNA crosslinks (21), and in the release of condensin complexes (22). Thus, Cdc48 is well-positioned to regulate protein sequestration pathways such as INQ following DNA damage detection. Other groups have noted the importance of SUMO to INQ formation (10), and we also find that reducing SUMOylation with *ubc9-ts* (SUMO E2) or *SIZ1* (SUMO E3) deletion reduces INQ localization of Hsh155-GFP in MMS-treated WT or *cdc48-4* cells (**Figure 3A** and **3B**). Indeed, mutation of the SUMO interacting motif of *CDC48* impairs its ability to rescue *cdc48-4* cells relative to WT *CDC48* (**Figure 3C**) (23). Since we used Hsh155-GFP as a marker for INQ we elected to directly test whether Hsh155 is SUMOylated using a Smt3-Hisx7 purification scheme (24). While SUMOylation of the Rfa1-GFP control was easily detectable upon MMS treatment, no detectable modification of Hsh155-GFP was observed with or without MMS (**Figure 3D)**. Thus, the identity of any SUMO targets important for regulating INQ will require additional study beyond the scope of this work. However, our findings again highlight the importance SUMO in regulating INQ.

**Figure 3.**
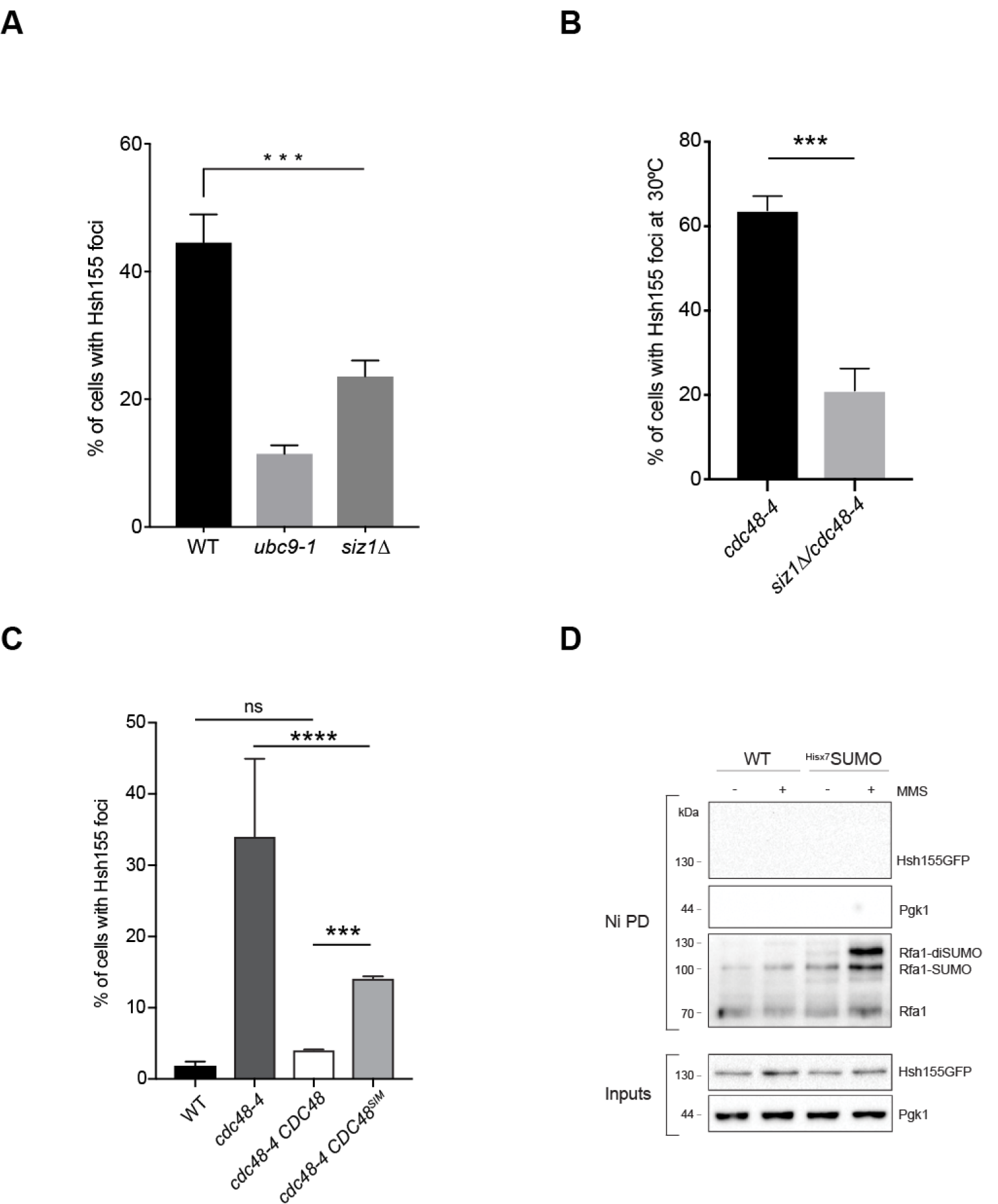
Testing potential roles of SUMO in Cdc48 induced Hsh155-GFP foci formation. (A) Hsh155-GFP foci induction by MMS in SUMO regulator mutant strains *ubc9-ts* and *siz1*Δ. (B) Induction of spontaneous Hsh155-GFP foci in *cdc48-4* is reduced by deletion of SUMO E3 ligase *SIZ1*. (C) Spontaneous Hsh155-GFP foci in *cdc48-4* complemented by WT or *cdc48-SIM* mutant bearing plasmids. For A-C quantifications: Error bars are mean± SEM, n= 3 with >50 cells each. Fisher test, Asterisks show *p*-value thresholds (**** p<0.0001; *** p<0.001 and non-significant (ns)). (D) SMT3-hisx7 pulldown assays in untreated or MMS treated cells. DNA damage induced SUMOylation of RFA1 in MMS (30) is shown as a positive control after Ni-NTA bead pulldown (Ni PD). No bands are detectable when blotting for Hsh155-GFP.

### Cdc48 regulates the stability of Hsh155 and assembly with its partners

If Cdc48 is important for the prevention or dissolution of INQ structures, then it could regulate the stability of INQ substrate proteins under stress. Consistent with this idea, Hsh155-GFP protein was more abundant in the *cdc48-4* mutant, and more stable in cycloheximide chase experiments, than it was in WT (**Figure 4A**). Since cycloheximide treatment can itself affect protein sequestration (7), we also monitored the impact of Cdc48 on Hsh155 lifetime with an orthogonal approach using *in vivo* tandem fluorescent timer (tFT) fusions (25). The ratio of GFP to mCherry fluorescence in Hsh155-tFT fusions is significantly red-shifted in *cdc48-4* cells indicating a longer relative protein lifetime (**Figure 4B** and **S1E**). These data are consistent with the INQ serving a protective role for substrate proteins as Hsh155 becomes both more abundant and aggregates in *cdc48-4* cells. Cdc48 could affect the stability of Hsh155 through direct protein-protein interactions. Indeed, we observed co-precipitation of Hsh155-GFP with Cdc48-TAP and via reciprocal pulldown (*i*.*e*. Hsh155-TAP/Cdc48-GFP) under both untreated and MMS treated conditions (**Figure 4C and D**).

**Figure 4.**
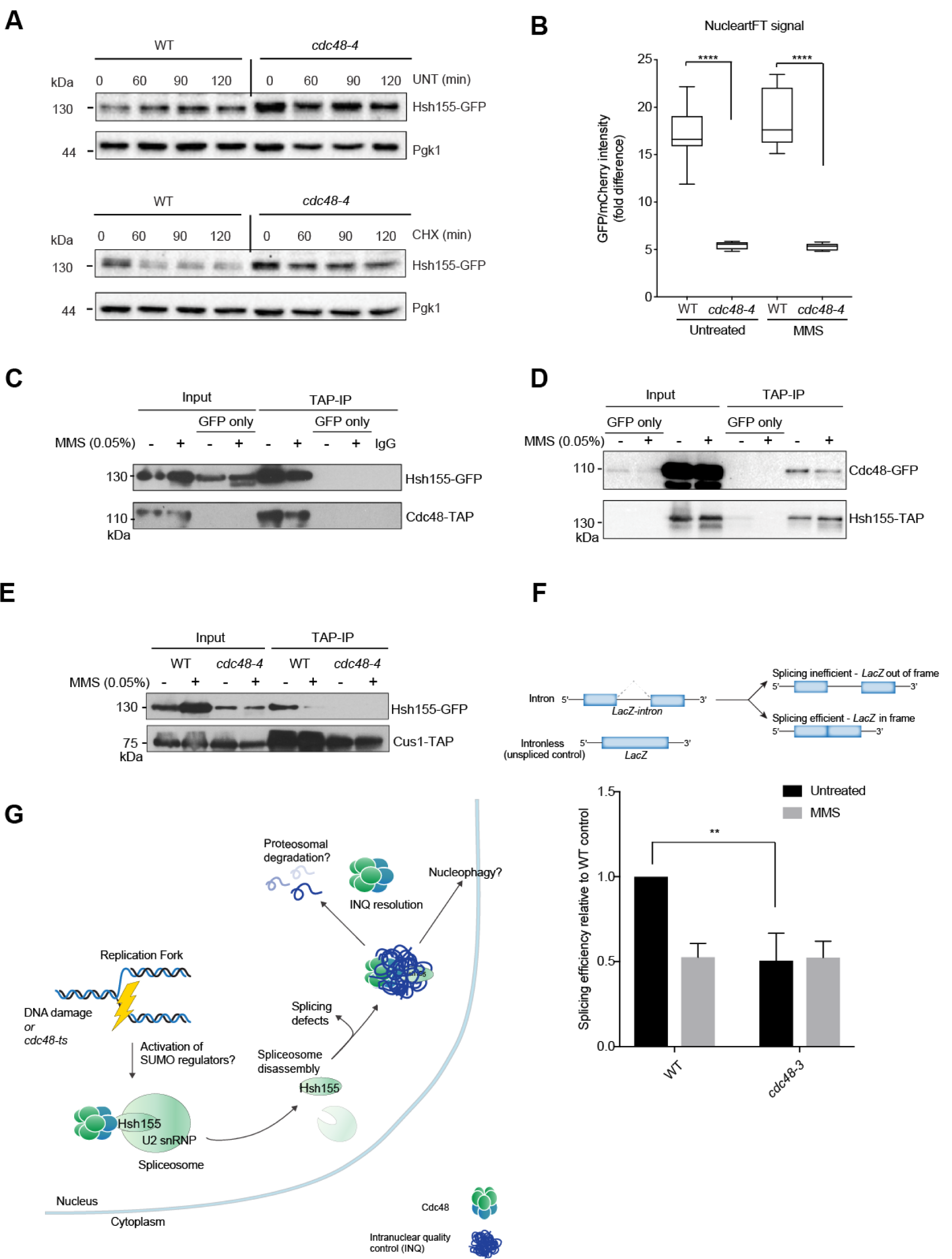
Direct effects of Cdc48 on Hsh155 stability and function. (A) Hsh155 protein levels ± CHX by anti-GFP western blotting relative to Pgk1 levels in WT and *cdc48-4* over time. CHX = cycloheximide, UNT = untreated. Shown is the representative blot from three independent experiments. (B) Tandem fluorescent timer (tFT) analysis of Hsh155-mCherry-GFP dual fusion lifetime in the indicated strains with or without MMS treatment. Ratios of GFP (fast maturing) and mCherry (slow maturing) fluorescence in the nucleus are shown. Quantification of fluorescent timer indicates the presence older proteins in the nucleus of *cdc48-4* compared to WT in both conditions. Mean ±SEM, three replicates, n ≥30 per replicate, Mann-Whitney test, asterisks show p-value threshold **** p<0.0001. (C and D) Physical interaction of Hsh155 and Cdc48 with or without MMS treatment by co-immunoprecipitation using Cdc48-TAP in C and Hsh155-TAP in D followed by western blotting. Control (GFP only) lanes are mock IPs from cells expressing Hsh155-GFP only. (E) Co-precipitation of Hsh155 with its partner Cus1 in the SF3B complex in *cdc48-4* is disrupted in both untreated and MMS conditions similar to MMS induced WT cells. (F) LacZ splicing reporter assay results in WT or *cdc48-3* treated with MMS. A schematic of the assay (above) shows the intronic versus intronless LacZ constructs that enable quantification of splicing efficiency. Quantification of relative LacZ splicing in untreated and MMS treated normalized to ‘no intron’ control in both WT and *cdc48-3* cells (below). Mean± SEM, Student t-test, Asterisks show *p*-value (**) p<0.01. N=5 with individual transformants in triplicates each. (G) Model of Cdc48 as a chaperone for Hsh155 involved in its quality control and sequestration at INQ sites (see main text). Cdc48 binds Hsh155 under normal conditions but its disruption or DNA damage leads to SF3B disassembly. Cdc48 may also play a role in INQ resolution or turnover, explaining why its loss increases INQ.

Hsh155 is a splicing factor that exists in a complex called SF3B with Hsh49 and Cus1. We previously found that Hsh155 disassembled from the SF3B complex upon MMS treatment (11). To assess the potential impact of Cdc48 on SF3B complex stability, we monitored pulldown of Hsh155-GFP in WT or *cdc48-4* cells with or without MMS treatment by co-immunoprecipation. Loss of *CDC48* function greatly diminished the presence of the Hsh155-Cus1 complex, even in untreated cells (**Figure 4E**). This suggests that Cdc48 may play a role in the integrity of the SF3B complex even in the absence of DNA damage signals. While *cdc48-4* grew too poorly to easily monitor splicing activity, the *cdc48-3* allele that showed an intermediate induction of Hsh155 foci (**Figure 2C**) did show reduced splicing flux in a LacZ-based splicing efficiency assay (**Figure 4F**) (26, 27).

### Context and perspective

Spatial control of PQC is a fundamental feature of stress responses in eukaryotic cells (28). Here we show that Cdc48, a key regulator of numerous cellular PQC activities, is also associated with INQ where it interacts, reduces the accumulation, or speeds the turnover, of the INQ substrate Hsh155 (**Figure 4G**). This defines a novel and complementary function for Cdc48 in nuclear protein quality control. Loss of SUMO-conjugating enzyme function, or E3 ligase function reduced INQ formation, consistent with a role for this modification in organized INQ structures. Considerably more work is needed to understand the regulation of INQ substrate protein localization and turnover. Our data specifically implicate Cdc48 in regulating the sequestration, complex formation and stability of the spliceosomal protein Hsh155 within the SF3B complex. We previously found that Hsh155 sequestration may regulate ribosomal protein gene expression through reduced splicing efficiency of ribosomal protein gene transcripts (11). In this regard, it is notable that Cdc48 is also a critical regulator of ribosomal quality control mechanisms and can sense ribosomal stress (29, 30). It is possible that these responses are coordinated during the DNA damage response. Regardless, our data define a new function for Cdc48 in regulating the deposition of proteins at the INQ and potentially in the regulation of splicing complexes. There is currently little evidence linking Cdc48 to splicing. Interestingly, a study of motor neuron transcriptome dynamics during iPS differentiation models of human VCP/Cdc48 mutations in amyotrophic lateral sclerosis showed that abnormal intron retention events were increased in the *Cdc48/VCP*-mutated cells (31). Thus, a role for Cdc48 in splicing across species is possible, and we hope that additional research will elucidate if and how Cdc48 may affect splicing.

## Materials and Methods

### Yeast strains, MMS treatments, and growth analyses

Yeast strains used in this study including database IDs and genotypes, primers and plasmids are listed in **Table S1**. All strains were in the s288c background and were grown under nutrient rich YPD media or synthetic media lacking amino acids (SC) unless otherwise indicated. Serial dilution assays and growth curve analyses were performed as described (11, 32). Briefly, cells with identical optical density (OD) were serially diluted ten-fold and spotted on the YPD plates with a 48-pin replica pinning manifold and incubated at indicated temperatures for 72 hours. Growth curves were analyzed in a Tecan M200 plate reader monitoring OD600nm every 30 minutes for 48 hours at 30 °C. For all MMS treatment conditions cells were exposed to 0.05% MMS (∼99% Sigma) in synthetic complete media for 2 hours unless otherwise noted. For growth curves, MMS was then washed out and replaced with fresh media.

### Live cell imaging, image acquisition, analysis and statistical methods

Imaging and subsequent analysis were performed as described (11). Log-phase yeast were mounted on slides pretreated with concanavalin A, in SC growth media. Immobilized cells were imaged using an Objective HCX PL APO 1.40 NA oil immersion 100× objective on an inverted Leica DMi8 microscope with a motorized DIC turret (for Differential Interference Contrast imaging) and a filter cube set for FITC/TRITC (for GFP and m-Cherry fluorescence imaging). The images were captured at room temperature using a scientific complementary metal oxide semiconductor (sCMOS) camera (ORCA Flash 4.0 V2; Hamamatsu) and collected using MetaMorph Premier acquisition software and post processed (including gamma adjustments, counting of cells with/without foci, foci intensity measurements for tandem fluorescent timer (tFT) experiments using ImageJ (National Institutes of Health). For all microscopy experiments, the significance of the differences was determined using Prism5 or higher (Graphpad). For foci intensity measurements in tFT experiments, samples were compared with t-tests or ANOVA; Graphpad performs F-tests for variance as part of this analysis. For comparisons of proportions, Fisher tests were used and p-values were Holm-Bonferroni corrected in the event of multiple comparisons. Sample sizes and specific statistical details for each image analysis are listed in the figure legends.

### Co-Immunoprecipitation and western blotting

Co-Immunoprecipitation was performed using yeast strains containing TAP tagged Cus1, Cdc48 and/or GFP tagged Hsh155 treated with or without MMS. TAP tagged proteins was captured using IgG sepharose fast flow beads (Sigma) and proceeded as described (33). Immunoblotting was carried out with mouse anti-GFP (ThermoFisher) and rabbit anti-TAP (ThermoFisher). For western blots, whole cell extracts were prepared by tricholoroacetic acid (TCA) extraction and blotted with mouse anti-GFP (ThermoFisher), rabbit anti-RFA (Agrisera) or mouse anti-PGK1 (Santa Cruz) antibodies as described (10).

### Protein stability time course

Overnight cultures of the indicated strains were diluted below OD 0.2 and allowed to progress into logarithmic phase before collection. The cells were treated with or without cycloheximide (CHX) at a concentration of 200 μg/ml for the indicated time (0-120 minutes). The final concentration of 2×10^7^ cells was collected by centrifugation and the whole cell extracts were prepared by TCA extraction followed by immunoblotting.

### Splicing efficiency assay

Splicing assay protocol was adapted and performed as described (26). All measurements were taken with three individual transformants per replicate, totaling five replicates. Cells were struck as a patch on SC-leucine, then replica plated to glycerol-lactate–containing SC medium without leucine (GGL-leu). Cells from each patch were inoculated in liquid GGL-leu media for 2 h at 34°C and induced with final 2% galactose for 1.5 h before treatment with final 0.05% MMS for 30 min. Cells were lysed and assayed for β-galactosidase assay using a Gal-Screen β-galactosidase reporter gene assay system for yeast or mammalian cells (Applied Biosystems) as per the manufacturer’s instructions and read with SpectraMax i3 (Molecular Devices). Relative light units were normalized to cell concentration as estimated by measuring OD600.

### Ni-NTA pulldown of SUMOylated proteins

SUMO conjugated proteins were detected by performing protein pulldowns under denaturing conditions as described (24). In brief, strains of interest were transformed with a plasmid containing His-tagged SUMO (Smt3-Hisx7) (Addgene) under the control of a copper inducible promoter. Logarithmically growing cells were harvested at OD_600nm_ 0.6-0.8 to a final concentration of 10^9^ cells by centrifugation (4000rpm, 5min at 4°C). Pelleted cells were washed and resuspended in 5ml of pre-chilled water. Cells were then lysed with 800µl of 1.85M NaOH/7.5% (v/v) 2-mercaptoethanol followed by 20 min incubation on ice. Protein precipitation was carried out by adding 800µl 55% TCA on ice for 20 minutes. Precipitates were collected by centrifugation (8000g, 20min, 4°C) and were resuspended in 1ml Buffer A (6M guanidine hydrochloride, 100mM sodium phosphate, 10mM Tris-HCl, pH 8.0) and incubated on a rotating block at room temperature for 1 hour to solubilise the precipitate completely. The resulting solution was then centrifuged (16,000g, 10min, 4°C) and supernatant was transferred to tubes containing 30µl precleared Ni-NTA agarose beads (Qiagen) in the presence of 0.05% Tween-20 and 15mM imidazole. These were incubated overnight on a rotating block at room temperature. Following the binding, the beads were washed twice with Buffer A containing 0.05% Tween-20 and four times with Buffer C (8M Urea, 100mM sodium phosphate, 10mM Tris-HCl, pH 6.3) containing 0.05% Tween-20. Beads were centrifuged (200g, 15s) and supernatant was completely removed. His-SUMO conjugates on the beads were eluted out by adding 30µl HU sample buffer and heating at 70°C for 10min. Resultant protein extracts were subjected to standard western blotting and probed for SUMO and other proteins of interest using antibodies as described above.

## Acknowledgements

P.C.S. is funded by a Natural Sciences and Engineering Research Council of Canada (NSERC) Discovery grant and Discovery Accelerator Supplement (RPGIN 2020-04360), and a Canadian Institutes of Health Research (CIHR) Project grant (MOP136982). P.C.S. is a CIHR New Investigator and Michael Smith Foundation for Health Research Scholar.

## Author Contributions

VM and PCS designed the project. VM, AK, YKJ, KW, AST collected the data and analyzed the data. VM, AK and PCS wrote the paper.

## Supplementary Information

**Figure S1.**
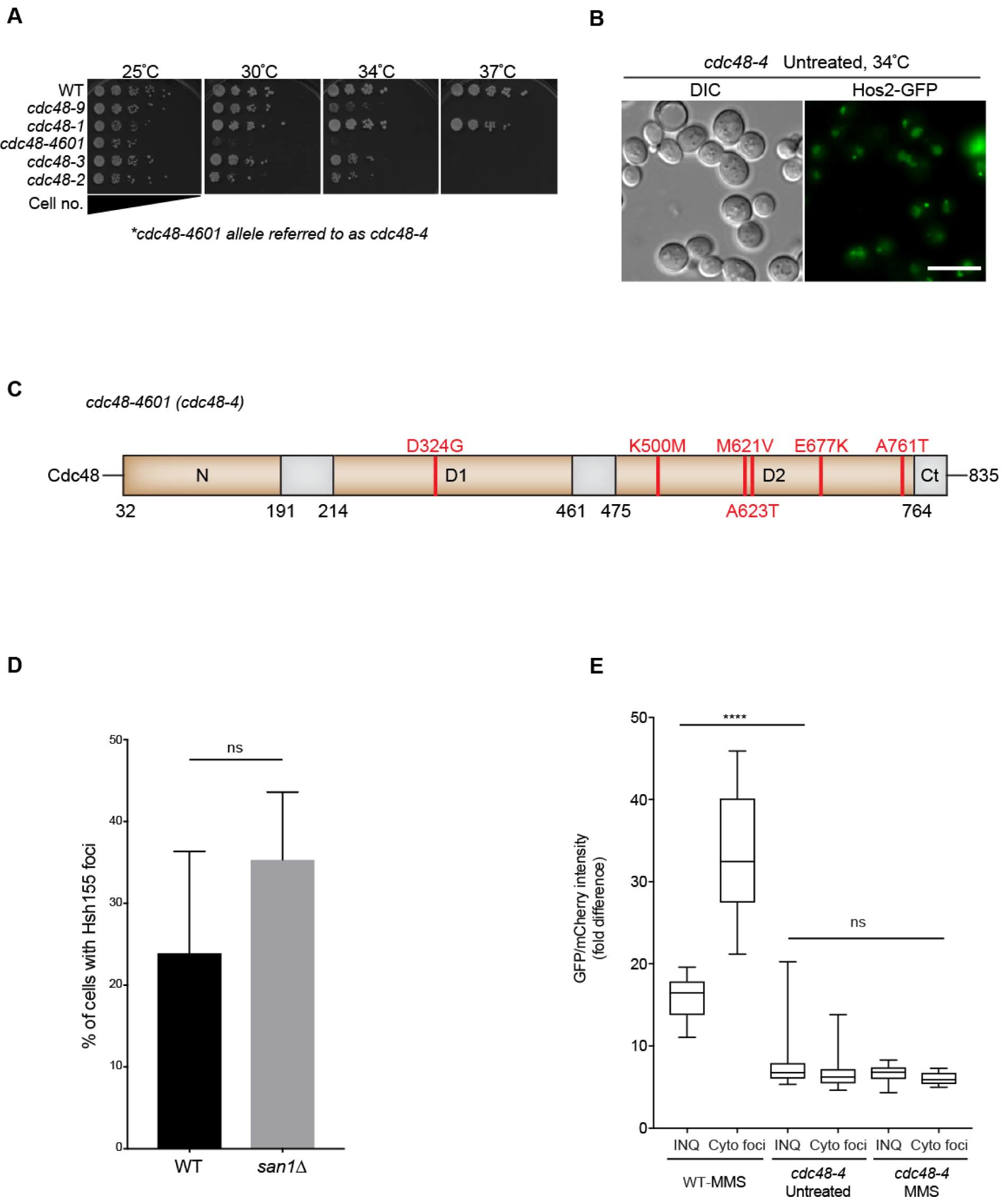
Characterization of the *cdc48-4* allele. (A) Spot dilution assay showing growth sensitivity of different *cdc48-ts* mutant alleles tested at the indicated temperatures. Equal ODs of the indicated strains were serially diluted and spotted on YPD at 25°C, 30°C, 34°C and 37°C. (B) Spontaneous Hos2-GFP foci formation in the *cdc48-4* mutant strain. (C) Schematic of Cdc48 protein domains (N-N-terminal, D1, D2 and Ct-C-terminal) with indicated *cdc48-4* coding mutations (red lines) mapped along the sequence. (D) Hsh155-GFP foci induction in WT or *san1*Δ strains treated with MMS. Error bars are mean± SEM, n= 3 with >50 cells each. Fisher test, non-significant (ns). (E) Tandem fluorescent timer ratio scoring of nuclear versus peripheral foci fluorescence in the indicated strains with or without MMS. Quantification of fluorescent timer indicates aggregation of older proteins in both INQ and CytoQ of *cdc48-4* compared to WT in both conditions. Mean± SEM, three replicates, n ≥ 30 per replicate, Mann-Whitney test, asterisks show *p*-value threshold **** p<0.0001; non-significant (ns).

**Table S1**. Yeast strains, primers, plasmids used in this study. See attached file.

